# A theory of the dynamics of DNA loop initiation in condensin/cohesin complexes

**DOI:** 10.1101/2021.11.10.468125

**Authors:** Bhavin S. Khatri

## Abstract

The structural maintenance of chromosome complexes exhibit the remarkable ability to actively extrude DNA, which has led to the appealing and popular “loop extrusion” model to explain one of the most important processes in biology: the compaction of chromatin during the cell cycle. A potential mechanism for the action of extrusion is the classic Brownian ratchet, which requires short DNA loops to overcome an initial enthalpic barrier to bending, before favoured entropic growth of longer loops. We present a simple model of the constrained dynamics of DNA loop formation based on a frictional worm like chain, where for circular loops of order, or smaller than the persistence length, internal friction to bending dominates solvent dynamics. Using Rayleigh’s dissipation function, we show how bending friction can be translated to simple one dimensional diffusion of the angle of the loop resulting in a Smoluchowski equation with a coordinate dependent diffusion constant. This interplay between Brownian motion, bending dissipation and geometry of loops leads to a qualitatively new phenomenon, where the friction vanishes for bends with an angle of exactly 180 degrees, due to a decoupling between changes in loop curvature and angle. Using this theory and given current parameter uncertainties, we tentatively predict mean first passage times of between 1 and 10 seconds, which is of order the cycle time of ATP, suggesting spontaneous looping could be sufficient to achieve efficient initiation of looping.

## Introduction

A fundamental feature of all life is the faithful replication and segregation of DNA into daughter cells during cell division, where one of the key steps is the condensation and separation of sister chromatids (*1*); the protein complexes condensin and cohesin that are part of the family of structural maintenance of chromosome (SMC) complexes and are highly conserved throughout all domains of life, are essential to this process and interact with DNA to regulate chromosome structure (*2, 3*). Condensin is known to play a key role in the compaction of chromosomes during mitosis, while cohesin acts to bind the sister chromatids together after DNA replication (*4*) and aids in chromosome alignment during cell division. Early electron micrographs showed the structure of mitotic chromosome consist of a central scaffold with loops of chromatin emanating from this central core (*5*). There is still much not understood about how these complexes act to achieve these compact structures and the separation of sister chromatids, but SMC complexes have been observed in vitro to exhibit the ability to actively extrude DNA into loops (*6–9*), which has formed the basis of the popular loop extrusion model of chromosome compaction: if a number of condensins bind to DNA and extrude loops, this will naturally lead to compaction of chromatin, with the condensins forming a backbone (*1, 10, 11*). Although, there are a number of empirical measurements that the loop extrusion model fails to capture (*12*) and there are a number of unanswered questions as to whether loop extrusion can in vivo lead to chromosome compaction, due to small stalling forces of motor activity (*6*) and how extrusion can occur in the face of nucleosome obstacles (*8*), SMC complexes show extrusion activity in vitro, which is poorly understood at a mechanistic level.

SMC complexes are essentially ring shaped protein trimers (*13*); they comprise a pair of coiled-coils which are bound to each other at their hinge domains, while the opposing end comprises the ATPase head domains that dimerise by ATP binding and are linked together by a mostly unstructured polypeptide called the kleisin subunit. Although, many of the molecular details remain to be firmly established and the exact path of DNA within the SMC complex is not known (known as topological, pseudo-topological and non-topological in the literature (*7, 14–16*)), a key feature for looping is that DNA is constrained or bound to the complex at two contact points (Fig.1), which enable the loop to grow. There are various models of how extrusion might occur, but the simplest is a classic Brownian rachet mechanism (*9*), where ATP hydrolysis causes unbinding of the head domains, allowing the DNA freedom to diffuse by Brownian motion with constrained electrostatic interactions with the head domains and additional protein domains known as HEAT repeat subunits,; the idea is that on re-binding of ATP if the loop has grown by diffusion then this motion has been ratcheted. Molecular motors powered by Brownian ratchet mechanisms work by relying on diffusion in some asymmetric potential, here for sufficiently long loops entropy will favour the growth of loops. However, for initially short loops the growth will be disfavoured by the enthalpy of bending. It is an open and important question whether Brownian motion would be sufficiently rapid for SMC complexes to initiate extrusion of DNA by this mechanism, or require additional force generation mechanisms to drive initial loop growth.

**Figure 1:**
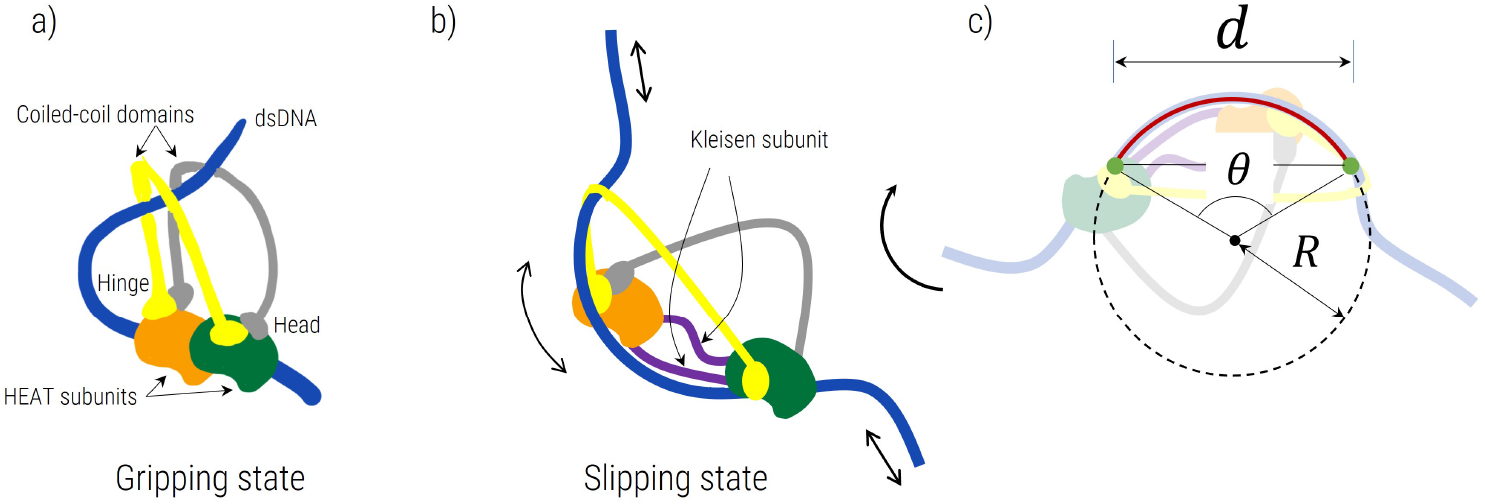
Diagram showing an idealisation of SMC complexes and DNA in gripping and slipping state (a & b) and c) how the looping of DNA is parameterised as an arc of a circle, where the *d* is the distance of the aperture constraining dsDNA and *R* is the radius and *θ* the angle of the loop. Note that although this diagram is based on the Brownian ratchet model (*9*) other models of extrusion can be represented by the same simple model shown in c) where the aperture *d* would correspond to a different distance.

From a physics perspective, the generic problem is one of the Brownian motion of a semi-flexible polymer loop through an aperture whose size is of order the persistence length. Although, there has been much theoretical (*17,18*) and empirical work (*19,20*) on the problem of DNA cyclisation, which studies the rate that short lengths of DNA find their ends, this is not directly relevant to looping within SMC complexes, as the constraints on DNA are different and the contour length of the loop can grow. These theoretical studies also ignore the role of internal friction, which is dynamical resistance to changing conformation due to internal energy barriers (*21–23*); it is well known from Kuhn’s theorem (*21, 24*) that for flexible polymers internal friction due to dihedral angle rotations are negligible for long polymers and the longest wavelengths of their Rouse modes, and conversely that for sufficiently short polymers they can be completely dominated by internal friction (*25*). The internal friction of proteins (*26–28*), unfolded polypeptides (*29*), polysaccharides (*23, 30*) and single stranded DNA (*31*) have all been measured empirically using a range of single molecule experimental techniques. A key message from these studies is that many dynamical mechanical processes in biology are far too slow to be explained if Stokes’ friction with the solvent was the only source of dissipation. In the case of semi-flexible or worm-like chain (WLC) polymers, measurements of stretched polypeptides (*29*) and ssDNA (*31*) show that internal friction increases with tension *F* with a power law ~ *F*^3/2^ predicted by a frictional worm-like chain (FWLC), which includes dissipation due to internal friction which opposes bending (*29*). The exact origin of bending friction is unclear, but will be related to steric constraints between complex molecular potentials giving rise to a local roughness that at a coarse-grained level gives an effective friction that opposes bending. Although, the bending friction constant of dsDNA has not been measured, we a priori expect the same behaviour as empirically determined for ssDNA and unfolded polypeptides.

In this paper, we address the fundamental theoretical question of Brownian loop growth by calculating the mean first passage time to reach a critical loop size such that entropy dominates, in the absence of any force generation mechanisms for loop growth. As we show, this is not a trivial diffusion problem, since for short loops of DNA we expect frictional resistance to bending (internal friction) within the polymer to dominate over Stokes’ friction with the solvent. We formulate the frictional worm like chain model to address this question and using Rayleigh’s dissipation function show that the constrained loop diffusion problem can be expressed as a 1 dimensional Smoluchowski equation in the loop angle with coordinate dependent friction. This analysis demonstrates a new physical phenomenon, which arises from an interplay between Brownian motion and loop geometry; we find this effective angular friction vanishes at exactly 180 degrees, since at this exact angle, infinitesimally, there is no change in curvature/bending of the loop and hence no bending friction. Further, given this phenomenon, we predict that even with relatively large initial angles of the loop, the mean first passage time to reach an entropically dominated loop is not significantly affected, and is dominated by a relatively small range of angles greater than 180 degrees. Given uncertainties of the exact parameter values – the internal friction to bending of double stranded DNA has not been measured – we make tentative predictions based on reasonable assumptions and estimate that loop initiation should take between 1 and 10 seconds. This time is of order the cycle time of ATP and suggests that initiation by the purely spontaneous Brownian looping described could be sufficient for efficient initiation of extrusion.

### Simple model of Brownian motion of DNA looping in SMC complexes

Fig.1 shows a diagram representing the simple model of how DNA looping can occur by diffusion in a SMC complex like condensin or cohesin, based on the Brownian ratchet model (*9*). The main assumptions of the simple model are that 1) DNA is bound to the protein complex at two points separated a distance *d* indicated with the small green circles, but free to slide frictionless through the point and with no imposed constraint in angle and 2) the conformation of the loop between the attachment points is an arc of a circle which has radius *R* (red line). After ATP hydrolysis and coiled-coil head disengagement, one of the binding points to DNA correspond to an electrostatic interaction with what is known as the head module (HEAT repeat subunit), which is bound to one of the coiled-coils, while the other in reality only provides a steric constraint on DNA by the same coiled-coil, but potentially constrained laterally by a kink in the coiled-coil referred to as the elbow; their separation gives the effective aperture of size *d* ≈ 25nm through which the loop can grow (*9, 32*). These assumptions are reasonable as more compact DNA conformations are likely to be entropically and energetically disfavoured and unlikely to contribute significantly to random paths that lead to looping through the complex. However, there are a number of uncertainties about the exact molecular details (*7, 14–16*), and there may be alternative paths DNA can take as it loops (topological vs pseudo-topological vs non-topological); a key element is that post-hydrolysis DNA has freedom to diffusively slide (slipping state) – constrained to different degrees by electrostatic binding to the head and hinge domains (via the HEAT subunits) – and upon ATP binding with the head domains and DNA, any Brownian loop growth is ratcheted (*9,32*). There are also very different models of how extrusion can occur within an SMC complex (*33, 34*), which broadly share the need for looping through an aperture, but where *d* would correspond to a different distance within the SMC complex. It should be further noted that this model does not include the initial search process and binding to DNA of the SMC complex, which means all estimates of mean first passage times below are lower bounds.

Given a circular loop we can write down the energy in terms of the radius *R* and the angle the loop subtends to it’s centre *θ*, given its persistence length 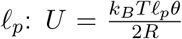. However, our simplification of the DNA strand being bound at both attachment points allows us to write down the radius in terms of the distance *d* and the angle *θ*, 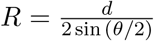, and so the energy of the loop as a function of *θ* only:

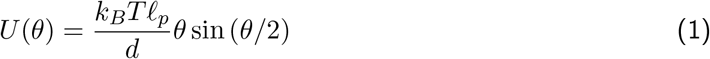

where increasing *θ* corresponds to increased progress of looping and a larger radius. We can now characterise the looping as a Brownian walk in the variable *θ* in a potential landscape given by *U* (*θ*), which is plotted in Fig.2a in units of *k_B_T* for *d* = 25nm and *ℓ_p_* = 50nm, where we see that maximum energy of a loop occurs at around *θ* ≈ 4 radians or 230°.

**Figure 2:**
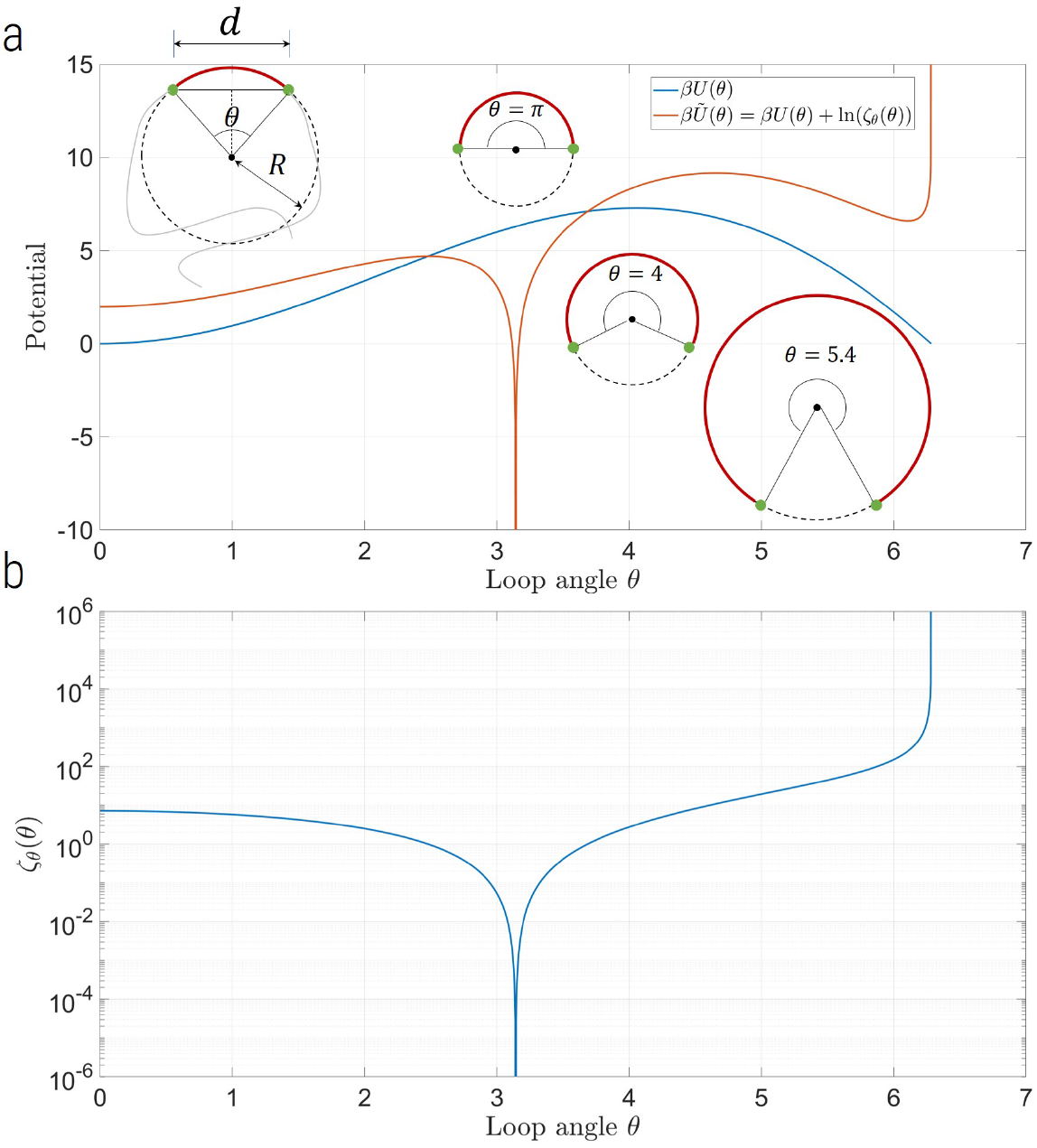
a) Potential energy *βU* (*θ*) (blue line) and the effective dynamical potential 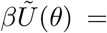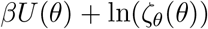 (red line) as a function of *θ*, where the latter incorporates the effect of internal friction in the dynamics of the loop. Note that the dynamic potential does not affect the Boltzmann distribution of angles which is still given by *U* (*θ*). The diagrams show the geometry of the DNA loop constrained to a distance *d* at two freely sliding attachment points, and also the relative loops sizes for different angles of *θ* = *π* (minimum friction), *θ* = 4 (maximum internal energy) and *θ* = 5.4, which corresponds to when the contour length of the loop *L* = 3*ℓ_p_* and when the loop will tend to adopt non-circular conformations. b) Effective angular friction of a loop *ζ_θ_*(*θ*) as a function of *θ*, showing vanishing friction at *θ* = *π* radians due the decoupling with changes in curvature at this angle.

The diffusion problem can be represented by the Smoluchowski equation:

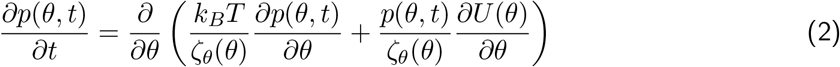

where *p*(*θ, t*) is the time-dependent probability density function of the angle *θ* as a function of time *t* and where the friction coefficient *ζ_θ_*(*θ*) is the effective coarse grained opposition to motion to changes in the angular velocity 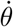, which as we will show is in general a function of the coordinate *θ*. The possible contributions to this effective angular friction are solvent dynamics in the nucleus and/or internal friction within the DNA, which dynamically opposes changes in conformation.

The natural model for the dynamics that encompasses both types of friction is the frictional WLC (*29, 35*), which is a model of semi-flexible polymer dynamics, which adds a term to the WLC model (*36–38*) that penalises changes in conformation that have rapid changes in curvature, as expressed by the following Langevin equation (*35*):

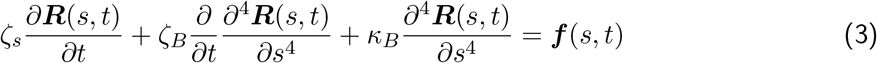

where *ζ_s_* ≈ 6*πη* is a solvent friction per unit length (*η* is the solvent viscosity), *ζ_B_* is the bending friction constant and *κ_B_* = *k_B_Tℓ_p_* is the bending elastic constant and *f* (*s, t*) is a spatially and temporally white noise term whose moments follow from the fluctuation dissipation theorem. A normal mode analysis shows the relaxation *τ_q_* of different modes has a mode- and length-dependent contribution to the relaxation which arises from solvent dynamics, whilst there is a mode- and length-independent contribution from internal frictional processes, since both internal friction and bending elasticity are coupled to curvature, they give the same dispersion relation with mode number:

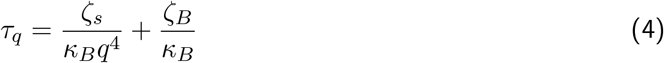

where *q* is the wavenumber of the mode. Comparing these two contributions for the relaxation of loops, whose contour length is of order the persistence length, then internal friction dominates the dynamics when *ζ_B_ ≫ ζ_s_*(*ℓ_p_/*2*π*)^4^; given *ℓ_p_* = 50nm and *ζ_s_* = 6*πη* ≈ 10^−5^*μ*g/(nm msec), we find *ζ_B_* 0.1*μ*gnm^3^/msec. If we treat the semi-flexible polymer as an viscoelastic rod with an effective internal friction per unit length *ζ_i_*, then simple arguments show that the bending friction should scale as *ζ_B_* = *ζ_i_r*^4^, with *r* is the radius of the rod (*35*). Although *ζ_B_* has not been measured for dsDNA, AFM experiments have estimated the bending friction for ssDNA as *ζ_B_* ≈ 11*μ*gnm^3^/msec (*31*) and *ζ_B_* = 0.3*μ*gnm^3^/msec for an unfolded polypeptide chain (*29*), which give an estimate of *ζ_i_* = 176*μ*g/msec/nm and *ζ_i_* = 187.5*μ*g/msec/nm, respectively. The consistency of these two estimates suggests that the local energy barriers or the local roughness of potentials that determine the effective internal viscosity of both ssDNA and unfolded polypeptides are similar, at least to order of magnitude, and so if we take *ζ_i_* ≈ 180*μ*g/msec/nm, then the internal friction bending constant of dsDNA will be *ζ_B_* ≈ 180*μ*gnm^3^/msec, which means for short loops of order a persistence length in contour length, internal friction will dominate and we only need consider this contribution to the Brownian motion of the loops. Alternatively, the double-stranded ladder structure of dsDNA may have a peculiar internal bending friction profile, which is not well approximated by a uniform viscoelastic rod, so at the other extreme we would expect dsDNA to have at least the internal friction of two strands of ssDNA acting in parallel, which would give twice the internal friction of ssDNA: i.e. *ζ_B_* ≈ 22*μ*gnm^3^/msec, which is still comfortably in the regime where internal friction dominates solvent dynamics.

Now as the loop angle performs a random walk the internal friction mediating this is related to the changes in curvature and so we need a way of calculating the coupling between these processes – in other words we want to translate a bending friction constant *ζ_B_* into an effective angular friction *ζ_θ_*. This can be done by using the Rayleigh dissipation function of the DNA loop; within a Langrangian formulation of mechanics, this in general is the dissipation rate of a system expressed in terms of the squared velocities and the friction constant of the system, and so by using the relationship between the curvature of the loop *κ* = 1*/R* = 2 sin (*θ/*2)*/d* and *θ*, we can derive an expression for the effective angular friction *ζ_θ_*. The Rayleigh dissipation function of the DNA strand is:

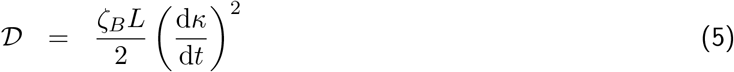

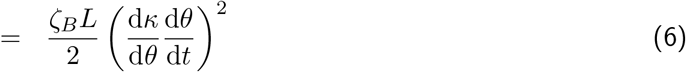

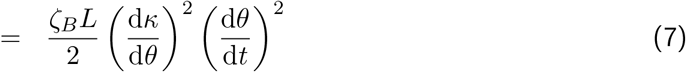

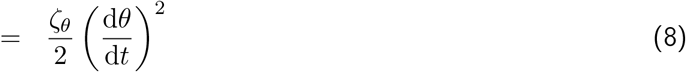

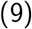

where *L* is the contour length of the loop between the points separated by *d*. This result means the effective angular friction of the loop is

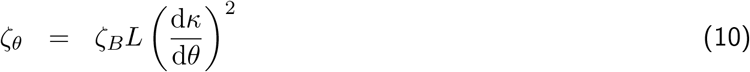

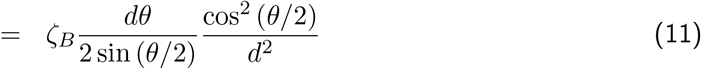

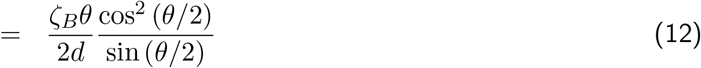

where we have used the fact that *L* = *Rθ*. We see that once translated to *θ* space the effective friction of the random walk has a very strong *θ* dependence, as plotted in Fig.2b, which shows in fact that the friction vanishes at exactly *θ* = *π* and diverges for *θ* = 2*π*. We can understand both these phenomenon in geometric terms; given the constraint that the loop emerges from an aperture of size *d*, when *θ* → *π* radians, small infinitesimal changes in the angle do not affect the radius of the loop, which means the curvature is unaffected and the friction must vanish, whilst the converse occurs as *θ* → 2*π* radians, where small changes in the angle lead to a very large changes in the size of the loop and so rapid changes in curvature, where in the exact limit, the radius of the loop must become infinite. Of course, a vanishing internal friction means that at angles close to 180 degrees Stokes’ friction will become dominant, which nonetheless we expect to make a negligible contribution to the mean first passage time and we ignore this in this treatment.

In practice, in the latter case once the contour length of the loop is much greater than the persistence length, the loop will adopt a very non-circular conformation before the divergence is reached. If we assume that once the contour length reaches a factor *α* larger than the persistence length, non-circular looping becomes dominant then these considerations give an approximate expression for the critical angle at which this occurs as

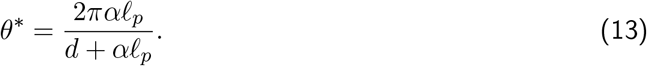

For *d* = 25nm, *ℓ_p_* = 50nm and *α* = 3, this gives *θ*^∗^ = 5.4 radians, or *θ* ≈ 310°, so this would predict that loops remain reasonably circular, before degenerating into a more random configuration.

### Mean first passage time for loop intiation

Given the Smoluchowski Eqn.2, using the flux over population method (*39*), it is possible to show that the mean first passage time (MFPT) to reach an angle *θ* is given by the following exact expression:

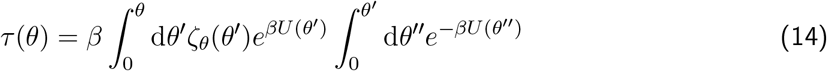

where *β* = 1*/k_B_T*. Although, various approximations can be obtained to evaluate this double integral, it is simple to numerically evaluate this instead, as shown in Fig.3 for different values of *θ*, where we take *ζ_B_* = 180*μ*gnm^3^/msec, the value we estimated for dsDNA; note that as *ζ_θ_* ~ *ζ_B_*, which can come out of this integral, such that the MFPT *τ* ~ *ζ_B_*, so that different assumptions about *ζ_B_* give a proportional scaling. We see that the MFPT increases rapidly for increasing angle *θ*, which diminishes as the loop approaches 180° at which there is an inflection point; this demonstrates the significant effect of the vanishing friction constant at *θ* = *π*. For angles greater than 180° the MFPT again rapidly increases, begins to plateaus for angles greater than *θ* = 5 radians, and then divergences for angles very close to *θ* = 2*π*, because of a diverging infinite friction, as observed above, although as discussed this divergence can be ignored, as in reality the chain will not be constrained to a circular conformation. For loop initiation dynamics in SMC complexes, the critical angle of interest is *θ* ≈ *θ*^∗^ ≈ 5.4, where the loop remains approximately circular. However, we see that on a log-scale the mean first passage time has already nearly plateaued for this angle; in other words the MFPT is strongly dominated by the time it takes to traverse a quite narrow range of angles, roughly between 4 and 5.4 radians or 230° and 310°.

**Figure 3:**
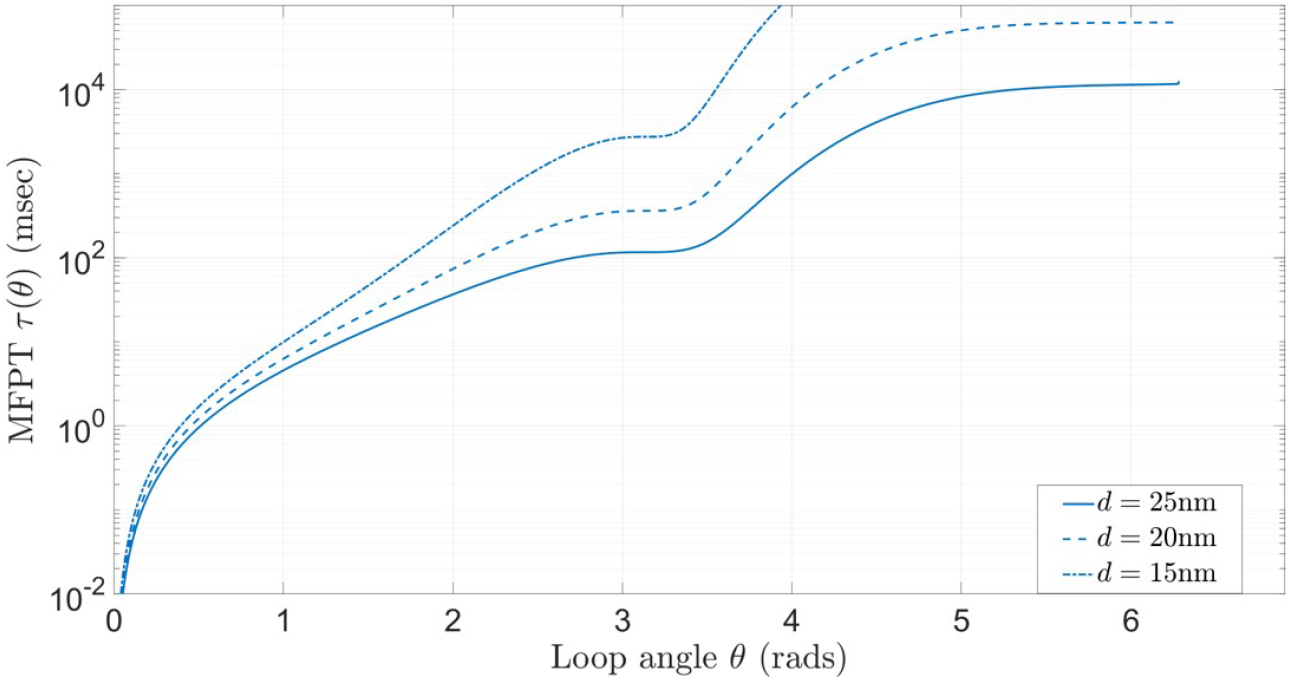
Mean first passage time *τ* (*θ*) (Eqn.14) for loop to reach angle *θ* assuming *ℓ_p_* = 50nm, *d* = 25nm, *ζ_B_* = 180*μ*gnm^3^/msec.

To understand this we note that the MFPT continues to increase rapidly even after the loop has passed the angle of maximum energy (*θ* ≈ 4 radians), which can be explained because of the still rapidly increasing internal friction of the loop; if we plot the effective dynamic potential 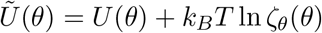, which appears in the integrand of Eqn.14 for the MFPT, then we see that it has a maximum at *θ* ≈ 4.5 radians and falls to more than *k_B_T* from the maximum at *θ* ≈ 4 radians and *θ* ≈ 5.4 radians. The angles which are within *k_B_T* of the maximum of the effective potential represents the regions over which diffusion is effectively flat, and relatively slow, due to the large internal friction at these angles.

## Discussion

The observation of loop extrusion within SMC complexes, like cohesin and condensin, suggests that such processes could be fundamental in the regulation of chromatin structure. Not least, it plays a central role in the simple “loop extrusion” model of chromosome compaction during mitosis, which is still an important and poorly understood fundamental biological process. Here we develop the simplest first model of how loops can grow to sufficient size by Brownian motion within SMC complexes, which is required for initiation of extrusion. We use this model of constrained Brownian motion of dsDNA, to calculate the mean first passage time (MFPT) to quantify how quickly such loops can form purely by the action of thermal Brownian motion, and so the degree to which loop initiation is dynamically constrained.

As we show small loops with contour length of order the persistence length are very likely dominated by internal friction to bending, rather than simple Stokes’ friction, which gives rise to a very different and to date little studied polymer dynamics problem, in general, and in particular with respect to understanding the dynamics of semiflexible polymers or worm-like chains like dsDNA. As the DNA chain is constrained to emerge from a aperture of fixed size, we can formulate the Brownian dynamics of looping using this frictional WLC model (*29, 35*), in terms of a 1D Brownian walk in the angle of the loop. From these considerations we see that bending friction and geometry interplay in a non-trivial way to determine the Brownian dynamics of loop growth, where friction vanishes at exactly 180° and the overall MFPT is dominated by a narrow range of angles approximately centred around a three-quarter circle (270°).

Evaluating the MFPT for *θ* = *θ*^∗^, we find *τ* ≈ 10, 000 msecs or 10secs and as the MFPT plateaus for angles approaching *θ > θ*^∗^ this estimate will be robust to order of magnitude given our assumption that dsDNA is a uniform viscoelastic rod with internal bending friction constant *ζ_B_* = 180*μ*gnm^3^/msec. Alternatively, if we assume the internal bending friction constant is at least the internal friction of two ssDNA strands in parallel, this would give *ζ_B_* = 22*μ*gnm^3^/msec (*31*), which is an order of magnitude smaller and thus giving an order of magnitude smaller estimate of *τ* ~ 1sec. The range of angles which dominate the MFPT are unchanged as changing *ζ_B_* just gives a constant vertical shift in the effective dynamical potential 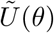. Empirical loop extrusion rates are of order ~ 0.1 to 1kbp/s (*6–9*), or ≈ 1 to 10*ℓ_p_*/s, and our result here suggests that the rate of initiation of loops is at least 1*/τ* = 0.1s^−1^ to 1s^−1^, since this calculation ignores the time to find and then bind DNA; although these two quantities need not agree, this suggests that loops are initiated at roughly the same rate, or slower than, the rate at which they are extruded, once initiated. However, more importantly the cycling time of ATP is of order ~ 1sec (*9*) and so this would suggest that if *ζ_B_* ≈ 180*μ*gnm^3^/msec corresponding to *τ* ~ 10secs, then typically Brownian motion alone would not quite be sufficient to generate a large enough loop in a single ATP cycle, although the probability would not be small, and may require a few cycles. If on the other hand *ζ_B_* ≈ 22*μ*gnm^3^/msec, which corresponds to *τ* ~ 1secs, then the probability of loop extrusion in a single cycle would be large.

There are other potential models of loop extrusion, such as the “tethered-inchworm” model (*33*) and the segment capture model (*34*), which postulate extrusion through the SMC complex in an unfolded conformation. These models do not discuss loop initiation and the simple model presented here is equally applicable, but with *d* > 25nm and a consequent expectation of a reduced mean first passage time for looping.

There are a number of simplifications in this first model of Brownian looping applied to loop initiation in SMC complexes. Firstly, it is clear that the binding of DNA to the hinge domain will not be frictionless or unrestricted in angle. In addition, there could potentially be torsional constraints on DNA on binding. Including some friction for sliding is relatively trivial and not affect the physical phenomenon due to the interplay of loop bending and internal friction discussed, but would tend to increase the quantitative estimates of MFPT. Regarding the constraints, given the electrostatic and non-specific nature of binding to DNA it is likely that once ATP is hydrolysed and the complex is in the slipping state any constraint on angle or torsion will be relatively weak. In any case, any such constraints would tend to be somewhat relaxed some distance along the contour of the loop and would likely give rise to a smaller effective aperture *d*, with a consequent increase in MFPT as shown in Fig.3. We have also ignored the out of plane entropy of the DNA loop, which will have some dependence on the contour length of the loop. However, as we have determined, the loop will maintain a roughly circular conformation up until the critical angle *θ*^∗^, with fluctuations about this mean conformation, such that entropic corrections will be logarithmic and relatively weak compared to the role of internal friction of the loops.

Finally, there is a large literature on the cyclisation dynamics of short lengths of dsDNA, which are of order a persistence length. The random walk problem of WLC like DNA finding its two ends is very different to the question considered here, since the constraints are different and the loop contour length can grow, although they both pertain to looping of DNA. Recent more careful experiments (*19, 20*) have shown that cyclisation times are very much longer (~ 1 minutes) than previously measured or predicted using the standard WLC. Despite the difference in constraints, the calculation of the regimes where internal friction vs solvent friction are important for contour lengths of dsDNA of order a persistence length are equally valid for the cyclisation problem and would suggest the standard WLC model would significantly underestimate cyclisation times due to its neglect of internal friction.

To summarise, we present a simple theory of Brownian loop growth in SMC complexes, where internal friction to bending of double stranded DNA couples to the changing geometry of the loop to determine the stochastic dynamics in a non-trivial way. For condensin/cohesin to act as a Brownian ratchet, loops must first grow to sufficient size to overcome the initial enthalpic barrier; the model remarkably predicts that friction for loop growth will vanish at 180° — with a corresponding plateau in the MFPT — and that the MFPT for the loop to grow large enough that entropy favours its growth is dominated loops that are roughly a three-quarter circle. This latter prediction conversely means that loop initiation is largely insensitive to the initial angle of the bend induced in DNA in the “gripping” state. Overall, we predict loop initiation times of order ~ 1s to 10s, which is of order the cycle time of ATP, suggesting loop initiation could initiate purely by Brownian motion without the aid of additional active force generation mechanisms.

## Acknowledgements

I thank Frank Uhlmann, Maxim Molodtsov and Tereza (Gerguri) Clarence for insightful discussions and comments on the manuscript.

## Data and materials availability

